# Modeling the impact of data sharing on variant classification

**DOI:** 10.1101/2021.06.21.449318

**Authors:** James Casaletto, Melissa Cline, Brian Shirts

**Affiliations:** Genomics Institute, University of California, Santa Cruz; Department of Laboratory Medicine and Pathology, University of Washington, Seattle

**Keywords:** Genetic variation, benign, pathogenic, classification, modeling

## Abstract

**Objective:** Many genetic variants are classified, but many more are designated as variants of uncertain significance (VUS). Patient data may provide sufficient evidence to classify VUS. Understanding how long it would take to accumulate sufficient patient data to classify VUS can inform many important decisions such as data sharing, disease management, and functional assay development.

**Materials and Methods:** Our software models accumulation of clinical data and their impact on variant interpretation to illustrate the time and probability for variants to be classified when clinical laboratories share evidence, when they silo evidence, and when they share only variant interpretations.

**Results:** Our models show that the probability of classifying a rare pathogenic variant with an allele frequency of 1/100,000 (1e-05) from less than 25% with no data sharing to nearly 80% after one year when labs share data, with nearly 100% classification after 5 years. Conversely, our models found that extremely rare (1/1,000,000 or 1e-06) variants have a low probability of classification using only clinical data.

**Discussion:** These results quantify the utility of data sharing and demonstrate the importance of alternative lines of evidence for the interpretation of rare variants. Understanding variant classification circumstances and timelines provides valuable insight for data owners, patients, and service providers. While our modeling parameters are based on assumptions of the rate of accumulation of clinical observations, users may experiment with the impact of these rates by downloading the software and rerunning the simulations with updated parameters.

**Conclusion:** The modeling software is available at https://github.com/BRCAChallenge/classification-timelines.

## OBJECTIVE

Genomic testing is now widely used for patients to determine if their genetics put them at increased risk of heritable disorders and to enable them to manage this risk clinically. For example, a patient with a known pathogenic variant in BRCA1 or BRCA2 should be screened more often for breast, ovarian, and pancreatic cancer. [1] Similarly, asymptomatic patients with familial cardiomyopathy might consider certain lifestyle changes such as losing weight, reducing stress, and sleeping well. [2]

The American College of Medical Genetics (ACMG) and the Association for Molecular Pathology (AMP) define qualitative, evidence-based guidelines for classifying genetic variants. Evidence for variant classification can come from many sources including clinical data, functional assays, and *in silico* predictors. Clinical data, typically derived through genetic testing reports, includes family history, co-segregation, co-occurrence, and *de novo* status. When sufficient evidence is present a variant curation expert panel (VCEP) may classify the variant as Likely Benign (LB), Benign (B), Likely Pathogenic (LP), or Pathogenic (P) using the ACMG/AMP rules for combining evidence. Variants with little or no evidence to support classification, called Variants of Uncertain Significance (VUS), create stress for patients and may lead to improper care. Because VUS do not yield medically actionable information, patients with VUS do not benefit from clinical management of their heritable disease risk. Ultimately, the significance of a variant remains uncertain until there is sufficient evidence to classify it. Although computational and functional predictions are helpful, some clinical data linking genotype and phenotype is usually needed to classify most variants. [3] However, there is no centrally available repository of clinical data that can be used for variant classification. Molecular testing laboratories and sequencing centers are the largest source of variant data. Many, but not all, clinical laboratories and sequencing centers actively share variant interpretations through ClinVar; however, they hold most of the clinical data they collect privately, due in large part to patient privacy and regulatory concerns. The shared interpretations for many genetic variants vary or even conflict between laboratories depending on the amount and nature of the evidence provided. [4]

One solution to these problems is for clinical laboratories to develop approaches to centrally share their clinical data associated with specific variants. Widespread sharing of variant pathogenicity evidence would lead to more rapid variant interpretation, greater scientific reproducibility, and novel discoveries. [4,5] Indeed, the National Institutes of Health recently mandated the sharing of all data for the research which it funds. [6]

While there is no question that data sharing would lead to expedited variant interpretation, and better patient outcomes by extension, under what circumstances is data sharing the most impactful? We have addressed this question by developing open source software to model the probability of variant classification over time under various forms of data sharing.

The output of our model not only quantifies the value of sharing variant data, the understanding of likely timelines and mechanisms of classification that this modeling illustrates could guide genetics organizations in prioritizing their efforts, inform strategies for functional assay development, improve variant classification guidelines, and enable healthcare providers to develop better strategies for managing specific patients with VUS. Further, the model serves as a platform for testing hypotheses on factors including the rates of gathering clinical evidence on the variant interpretation timeline. While we have informed the model with data on the frequency of observing various forms of clinical evidence according to the scientific literature and our own clinical experience, these factors are modeling parameters that can be modified easily as new evidence emerges, or to test the impact of clinical assumptions on the variant classification rate.

## MATERIALS AND METHODS

This section outlines a statistical model that combines clinical information from multiple sequencing centers to create an aggregate, pooled center so that VUS may be classified faster.

### Combining multiple forms of variant classification evidence

The evidence that the ACMG/AMP uses to classify variants encompasses several sources of data, including the type of variant (e.g. nonsense or frameshift), *in vitro* functional studies, *in trans* co-occurrence with a pathogenic variant, co-segregation in family members, allele frequency, and *in silico* predictions. They are divided into four levels of strength: “Supporting” (or “Predictive”), “Moderate”, “Strong”, and “Very Strong”. For example, PP1, which represents co-segregation of the disease with multiple family members, is considered “Supporting” evidence for a Pathogenic interpretation. Another form of evidence called BS4 represents the lack of segregation of the disease with the variant in affected family members. The BS4 evidence is considered “Strong” evidence for a Benign interpretation. [7] Tavtigian et al [8] showed that the rule-based ACMG/AMP guidelines can be modeled as a quantitative Bayesian classification framework. Specifically, the ACMG/AMP classification criteria were translated into a naive Bayes classifier, assuming the four levels of evidence and exponentially scaled odds of pathogenicity. While the ACMG/AMP guidelines define rules for the combinations of evidence which lead to variant classifications, the Bayesian framework assigns points to each form of evidence, where these points are summed and compared to thresholds to determine the variant’s pathogenicity. We leverage this Bayesian framework to model calculating odds of pathogenicity conditioned on the presence of one or more pieces of evidence for a given variant. For more detail regarding the combination of evidence, see Equations S1-S3 in the supplementary material.

For each variant, our model calculates two odds of pathogenicity: the odds of a VUS being benign and the odds of a VUS being pathogenic, both of which are conditioned on statistically sampled evidence.

### Selecting categories of variant evidence for model

Some sources of variant classification evidence are not impacted by data sharing, such as *in silico* prediction scores and functional assay scores. We will not use those categories of evidence in our model so we can specifically quantify the unique contribution of cumulative clinical data to variant interpretation.

Several sources of clinical case and family information will contribute to variant classification over time. As clinical databases grow and data is shared more effectively across institutions, more variants will be classified. Increased clinical information is the major source for variant reclassification as well. [9] We selected the following categories of clinical pathogenic evidence for our model

- *de novo* variants without paternity and maternity confirmation (PM6)
- co-segregation in family members affected with the disease (PP1)
- *de novo* variants with both paternity and maternity confirmed (PS2)

Similarly, we selected the following categories of benign evidence criteria that relate to clinical information

- *in trans* co-occurrence with a known pathogenic variant (BP2)
- disease with an alternate molecular basis (BP5)
- lack of segregation in affected family members (BS4)

The more evidence that is gathered over time, the sooner and more likely a VUS will be classified. However, not all the evidence that is gathered over this time will be concordant. [10] Patients who have a pathogenic variant may occasionally present evidence from one or more benign categories, for example, lack of segregation in affected family members due to disease heterogeneity. This presentation of conflicting evidence for a given variant occurs at a low, non-zero frequency. Therefore, we use a combination of pathogenic and benign evidence in the classification of every VUS.

### Parameters affecting clinical observations

To model the accumulation of clinical evidence, we defined certain modeling parameters according to the literature and to our own clinical experience. While the values that we have assigned to these parameters constitute well-informed assumptions, these values can be modified to test hypotheses, or as new knowledge emerges over time. In our software, these parameters are encapsulated in a single JSON file, so rerunning the model with revised parameter values requires modifying only one file.

#### Frequency distribution for evidence

Tavtigian et al calculated the corresponding odds of pathogenicity for each category of evidence and showed that the numerical-based odds are consistent with the rule-based ACMG/AMP guidelines for combining evidence.

**Table 1:** Odds and frequency estimates per ACMG/AMP category per evidence category.

| **Evidence category** | **Estimated benign evidence frequency** | **Estimated pathogenic evidence frequency** | **Pathogenicity odds of evidence** |
| --- | --- | --- | --- |
| **PS2** | 0.0015 | 0.003 | 18.7 |
| **PM6** | 0.0035 | 0.007 | 4.3 |
| **PP1** | 0.01 | 0.23 | 2.08 |
| **BS4** | 0.1 | 0.0001 | 1/18.7 |
| **BP5** | 0.07 | 0.0001 | 1/2.08 |
| **BP2** | 1.0 * frequency | 0.005 * frequency | 1/2.08 |

Tavtigian et al calculated the corresponding odds of pathogenicity for each category of evidence and showed that the numerical-based odds are consistent with the rule-based ACMG/AMP guidelines for combining evidence. Specifically, they determined that, for pathogenic evidence, the odds for “Strong” evidence is 18.7, for “Moderate” is 4.3, and for “Supporting” is 2.08. For benign evidence, the odds for “Strong” evidence is 1/18.7, for “Moderate” it’s 1/4.3, and for “Supporting” it’s 1/18.7. To estimate the frequencies of observing each form of clinical evidence, we examined the scientific literature. [11-14] Table 1 depicts the odds and estimated frequencies for the ACMG/AMP evidence categories that correspond only to clinical evidence. There may be pathogenic evidence observed for benign variants and benign evidence observed for pathogenic variants, though such observations generally occur at a low rate. For example, the frequency of BP2 for pathogenic variants is very unusual, except in tumors or in the case of rare diseases such as Fanconi anemia. Conversely, we assume that the frequency of BP2 evidence for benign variants is quite common and so occurs at the same rate as the variant itself.

#### Thresholds for odds of pathogenicity

Tavtigian et al defined four threshold ranges for the odds of pathogenicity for each of the four ACMG/AMP variant classifications (Benign, Likely Benign, Likely Pathogenic, Pathogenic).

**Table 2:**
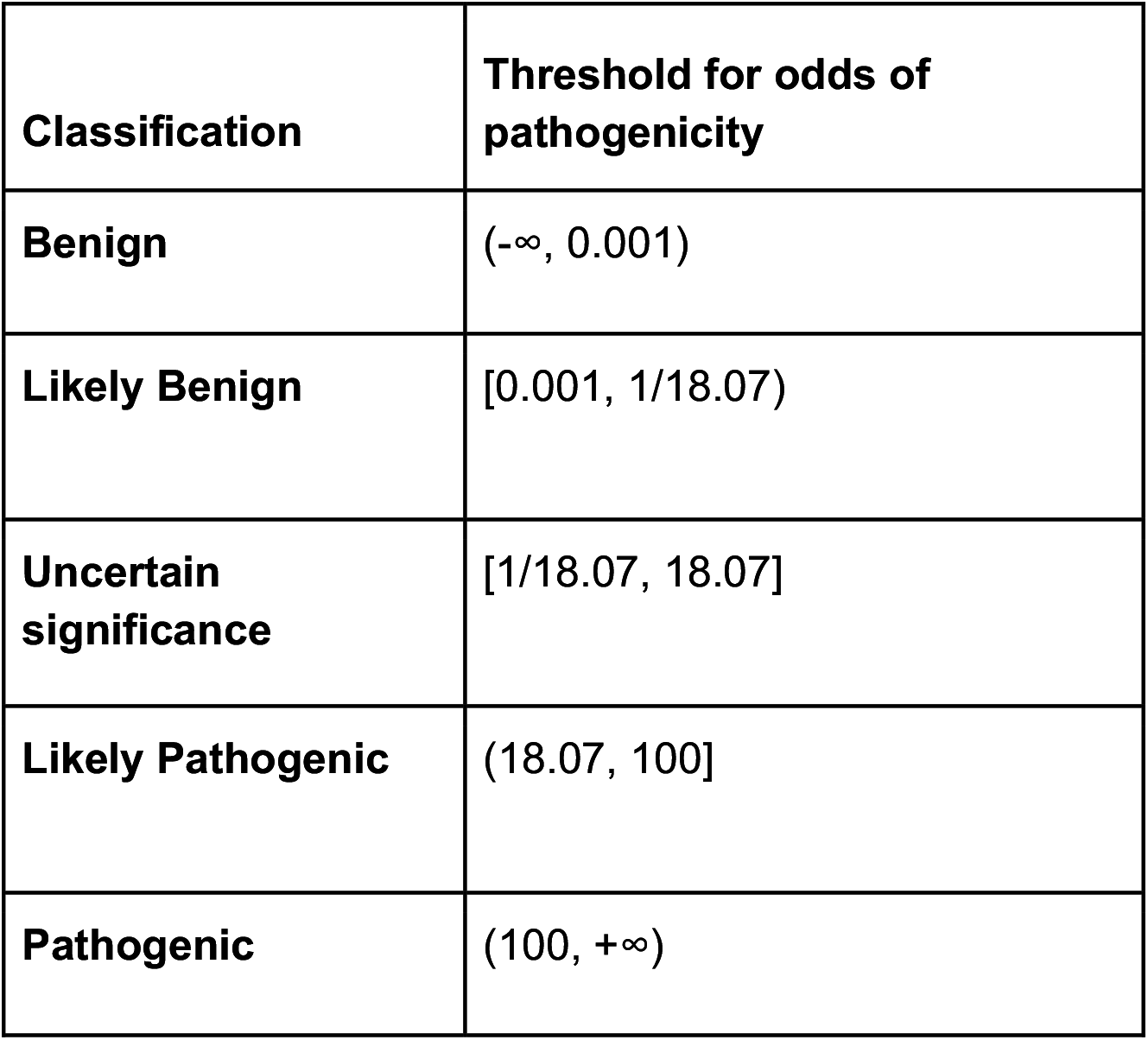
Odds of pathogenicity per classification.

These thresholds correspond to the values from Tables 1 and are consistent with the ACMP/AMP rules for combining evidence. For example, having one piece of strong evidence (e.g. BS4) and one piece of supporting evidence (e.g. BP2) is sufficient to classify a variant as “Likely Benign”.

#### Data from participating sequencing centers

For generating simulated clinical data, we define three categories of sequencing centers: small, medium, and large as shown in Table 3.

**Table 3:** Current number of tests in database and testing rate per center type.

| **Center type** | **Current number of patients tested** | **Tests per year** |
| --- | --- | --- |
| **Small** | 15,000 | 3,000 |
| **Medium** | 150,000 | 30,000 |
| **Large** | 1,000,000 | 450,000 |

We estimated the large database size and testing rate from the online publications of relatively large sequencing labs, [15, 16] and we estimated the small database size and testing rate based on our own experience at the University of Washington Department of Laboratory Medicine (a relatively small laboratory). We estimated the medium database size and testing rate by interpolating between the large and small database values.

#### Ascertainment bias

Healthy people from healthy families are underrepresented in many forms of genetic testing. [17] Accordingly, patients with pathogenic variants are observed (or ascertained) more often than those with benign variants, and the forms of evidence that support a pathogenic interpretation accumulate more quickly. How much more likely a person is to present pathogenic evidence than benign evidence is captured in our model as a real-valued constant. We conservatively estimated this term to be 2 based on our experience at the University of Washington Department of Laboratory Medicine.

#### Prior odds of pathogenicity

The Bayesian prior odds of a variant’s pathogenicity represents all other criteria that are not clinical and do not change much, if at all, over time. For this implementation, we sampled a random value from a uniform distribution between 1/18.07 and 18.07 which is the lower and upper bound of the odds of pathogenicity for VUS.

### Implementing the simulation

Our statistical model contains one variable: the allele frequency of the VUS of interest. Parameters of the simulation software include the number and types of each of the participating sequencing centers and the number of years for which to run the simulation. Because the variant is of uncertain significance, we gather evidence for both benign and pathogenic classifications simultaneously.

For the first year of our simulation, all the evidence that is assumed to be currently present at each of the individual testing centers is initialized and aggregated. We use the Poisson distribution sampling method when determining how many times the variant is observed, given the VUS frequency. For each year in the simulation, we generate new observations for variants assumed to be benign and assumed to be pathogenic at each sequencing center. We aggregate those observations across participating centers into a single collection to simulate the sharing of data.

We ran simulations as described above 1,000 times to simultaneously generate data points for VUS which occur at the rate of one in every 100,000 people (1e-05), combining data from 10 small centers, 7 medium centers, and 3 large centers generated over 5 years. We then ran similar simulations (1,000 times) for a VUS of frequency 1e-06 (one in every 1,000,000 people) in the same grouping of centers.

We created histograms and scatter plots that show the distribution and progression of the evidence over time. For each year, we plot the probability that each center classifies the variants individually using siloed data or if they collectively pool their data. We calculated the probability of a variant being classified at any sequencing center using the inclusion-exclusion principle in probability [18] assuming all centers would share all variant interpretations. This is a conservative estimate: not all sequencing centers share all their variant interpretations. We performed a sensitivity analysis to show the impact that each of the evidence types has on the probability of being either benign or pathogenic.

## RESULTS

In this section, we discuss the results of our simulation with variants over the course of 5 years at 20 participating sequencing centers. We first examine the histograms of the evidence for pathogenic and benign variants after 5 years of observations. Second, we examine the trajectory of evidence over the course of 5 years in scatter plots. Third, we examine the probability scatter plots over the course of 5 years. Fourth, we analyze the sensitivity of our results with respect to each type of evidence. These four sets of results were generated using a variant of 1e-05 frequency. Last, we examine the probability scatter plots over the course of 5 years for a 1e-06 (one-in-a-million) variant.

### Histogram plots of variants occurring at 1e-05 frequency

The distribution of evidence gathered individually and combined across all sequencing centers is plotted in Figure 1. As expected, increasing the number of classification data points for the many different variants results in wider Gaussian distributions that increasingly separate from the null assumption of no clinical evidence. More evidence provides more certainty in classifications as evidence exceeds the classification thresholds for an increasing number of variants.

**Figure 1:**
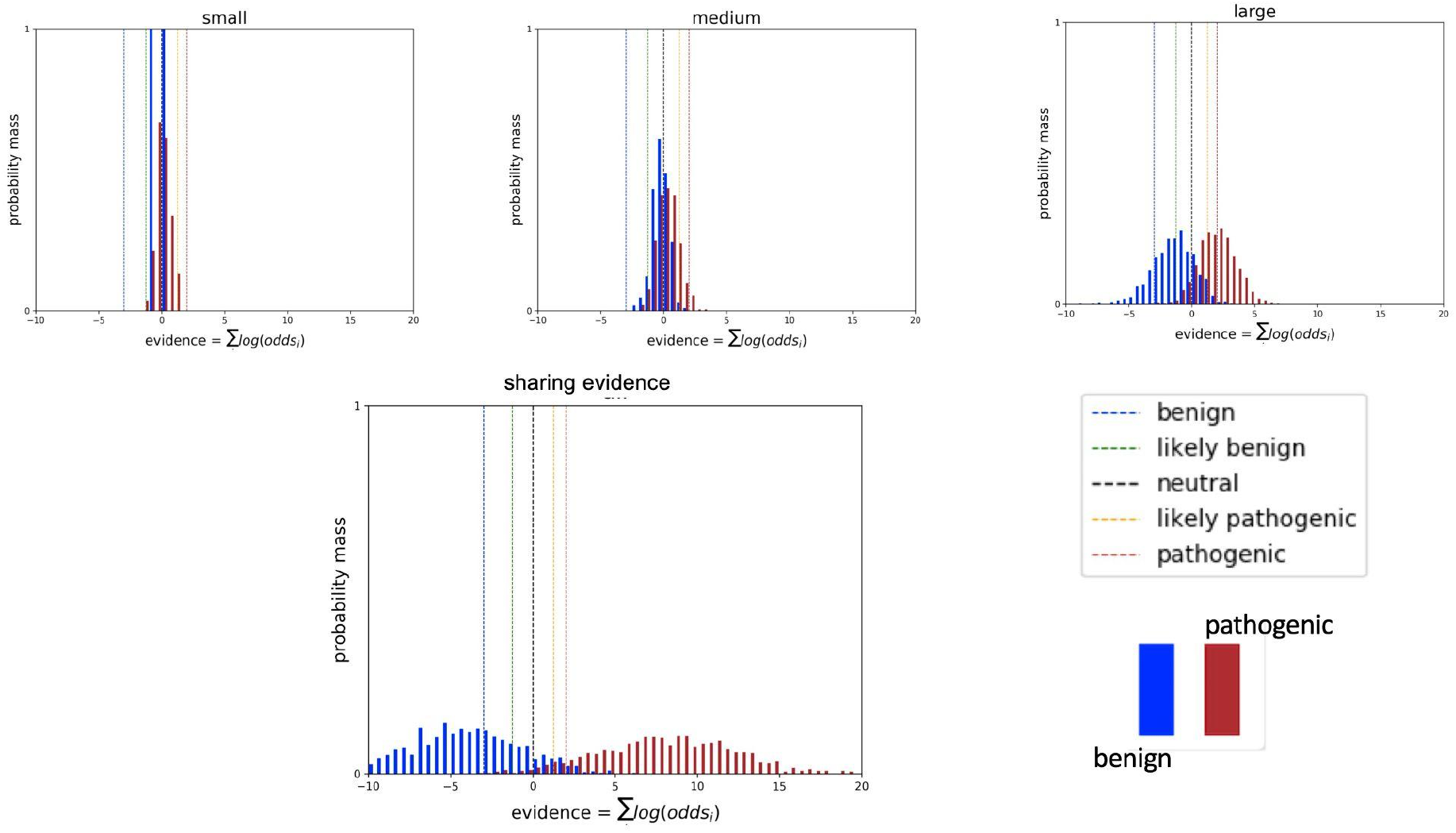
Histograms of cumulative log odds for classifying each of 1000 simulated variants present at a 1e-05 frequency in the population. Classification thresholds are demarcated as vertical hash lines. Benign variants are in blue and pathogenic variants in red.

### Trajectories of evidence for variants at 1e-05 frequency

The classification trajectory for individual variants can vary depending on which observations are made and when those are made. Although data accumulation increases the likelihood of classification and the likelihood of correct classification for variants as a group, evidence for individual variants may rise and fall. Figure 2 plots a subset of 20 classification trajectories (10 benign and 10 pathogenic) at a small, medium, and large sequencing center as compared to the combined data across all sequencing centers assumed to be sharing evidence. Trajectories in these scatter plots mimic real-world phenomena: variants may accumulate contradictory evidence; and long time periods may pass with insufficient evidence.

**Figure 2:**
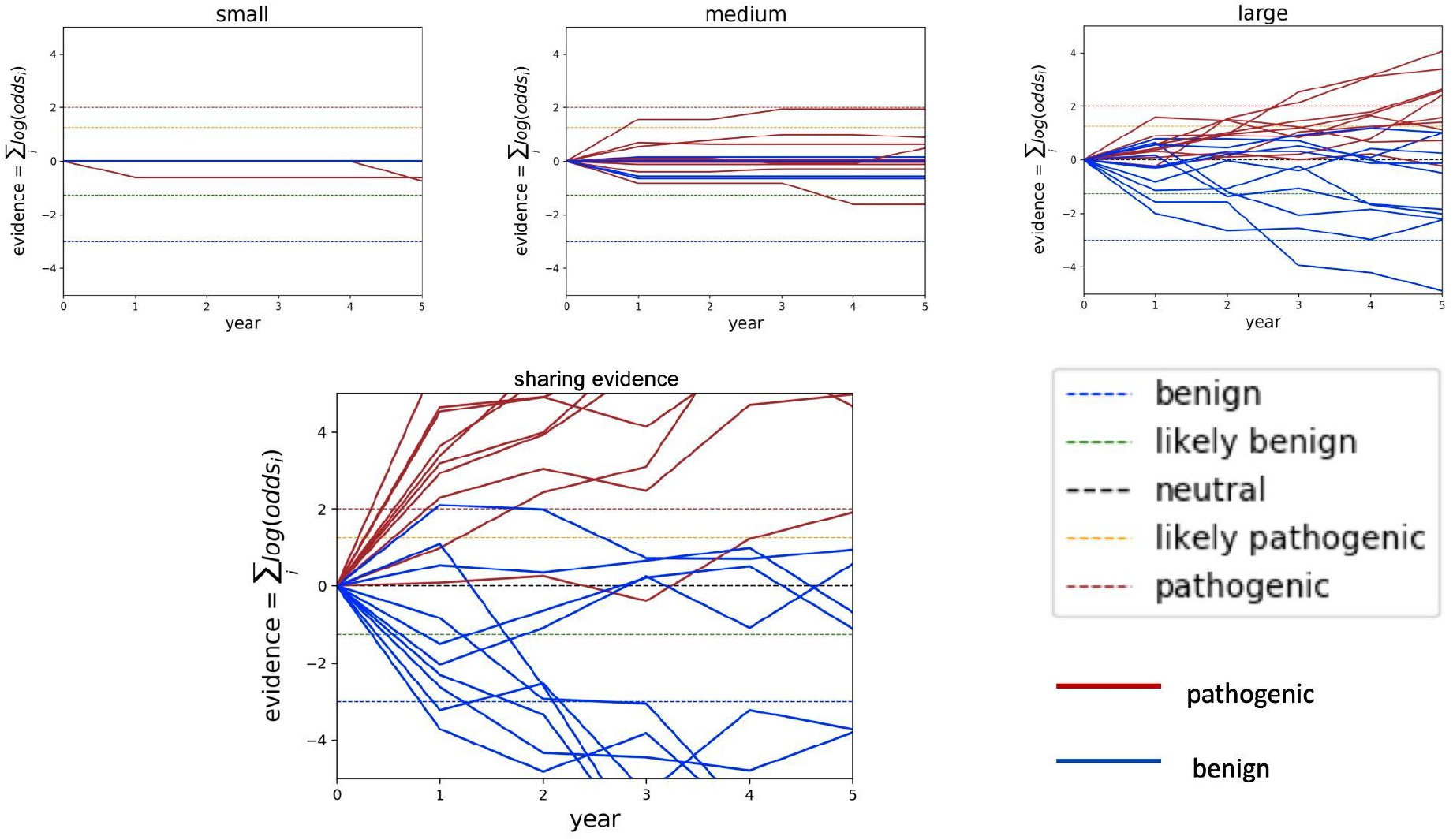
Classification trajectories for 20 randomly selected variants at 1e-05 frequency in the population. Classification thresholds are demarcated as horizontal hash lines in the timeline plots. Benign variants are in blue and pathogenic variants in red.

### Probabilities of classifying variants at 1e-05 frequency

Figure 3 shows the probability of classifying a variant which occurs at 1e-05 frequency in the population over the course of 5 years under different sharing paradigms. We show a small, medium, and large sequencing center not sharing anything as compared to two forms of sharing: centers sharing their all their variant interpretations but none of their clinical data (labeled “sharing classifications”); and centers sharing all their clinical data (labeled “sharing evidence”). From these graphs, we see that any data sharing increases the likelihood of variant classification and that sharing evidence rather than sharing classifications makes variant interpretation more certain by moving “Likely Benign” variants into the “Benign” classification and similarly moving “Likely Pathogenic” variants into the “Pathogenic” classification. Moreover, sharing evidence rather than sharing classifications reduces the amount of time required to classify variants.

**Figure 3:**
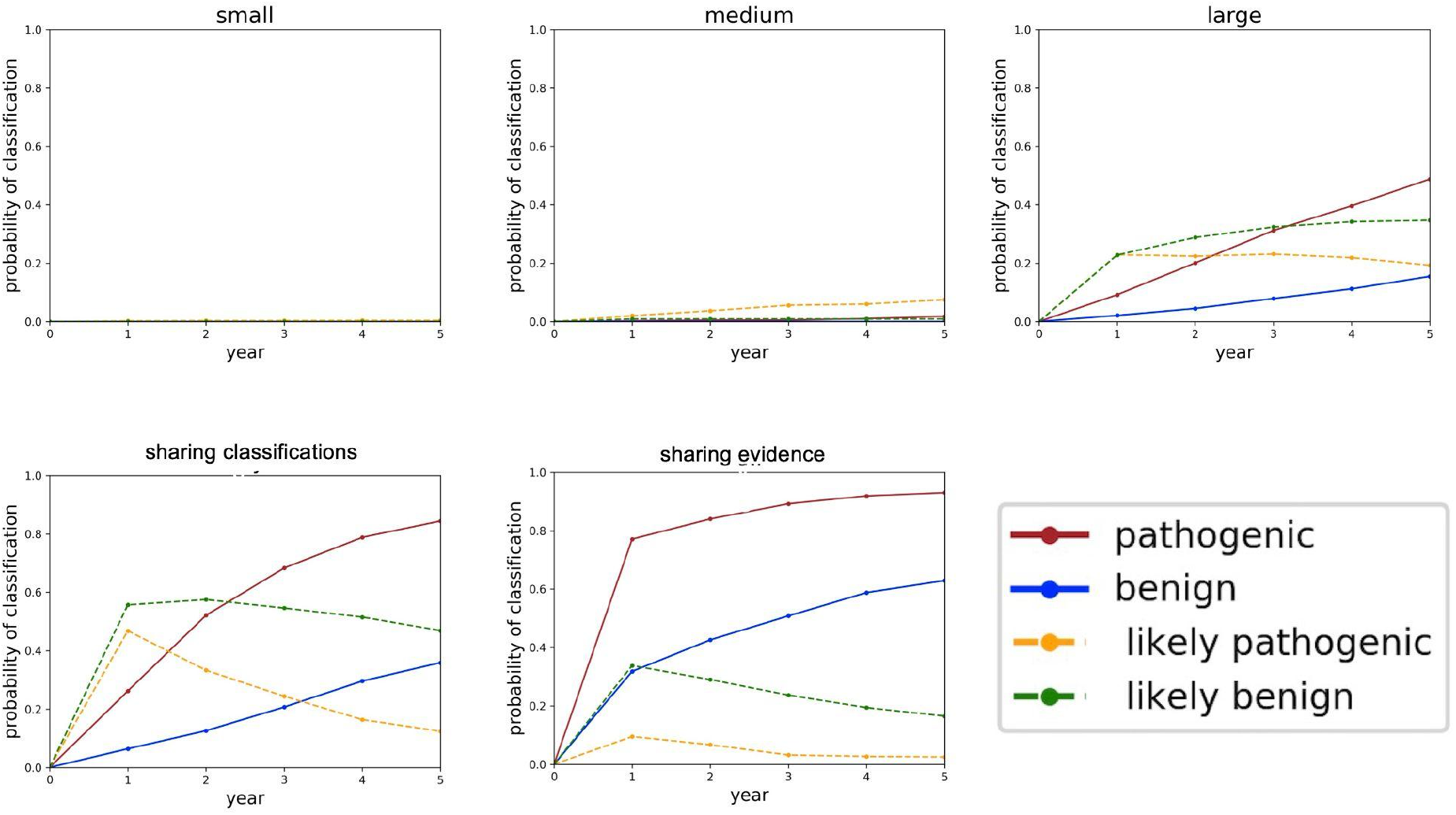
Classification probabilities over the course of 5 years. The y-axis of these plots is the probability of classifying the variant, converted from the aggregated likelihoods of pathogenicity generated in the simulations. After the first year in the sharing-evidence model, some of the variants which were LB shifted to B, and similarly some of the variants which were LP shifted to P.

In the supplement, we explore changing the distribution of sequencing centers and the number of years sharing data. In supplementary Figure S1, we see that after 20 years of data sharing, almost all benign and pathogenic variants are classified using clinical data alone. In supplementary Figure S7, we see that reducing the number of participating sequencing centers from 10, 7, and 3 (small, medium, and large) to 5, 3, and 1 significantly reduces the probability of classifying variants using clinical data alone.

### Sensitivity analysis for variants at 1e-05 frequency

We estimated conservative confidence intervals around the evidence observation frequencies defined in our model to determine how sensitive the probabilities of classification were to each type of ACMG/AMP evidence. We held all other parameters constant (equal to their expected values) while changing one frequency at a time to the low and high value in their respective interval to determine how sensitive the model is to changes in the frequencies observing different types of clinical data. Based on the assumptions of our experiments, classification of pathogenic variants is most sensitive to BS4 and PP1 evidence criteria (Figure 4a). Classification of benign variants is most sensitive to BS4 and PP1 evidence criteria (Figure 4b). Classifications were not affected by the change in BP2 evidence frequencies for either Benign or Pathogenic variants. Pathogenic variant classification was not affected by changes in BP5 evidence frequency.

**Figure 4:**
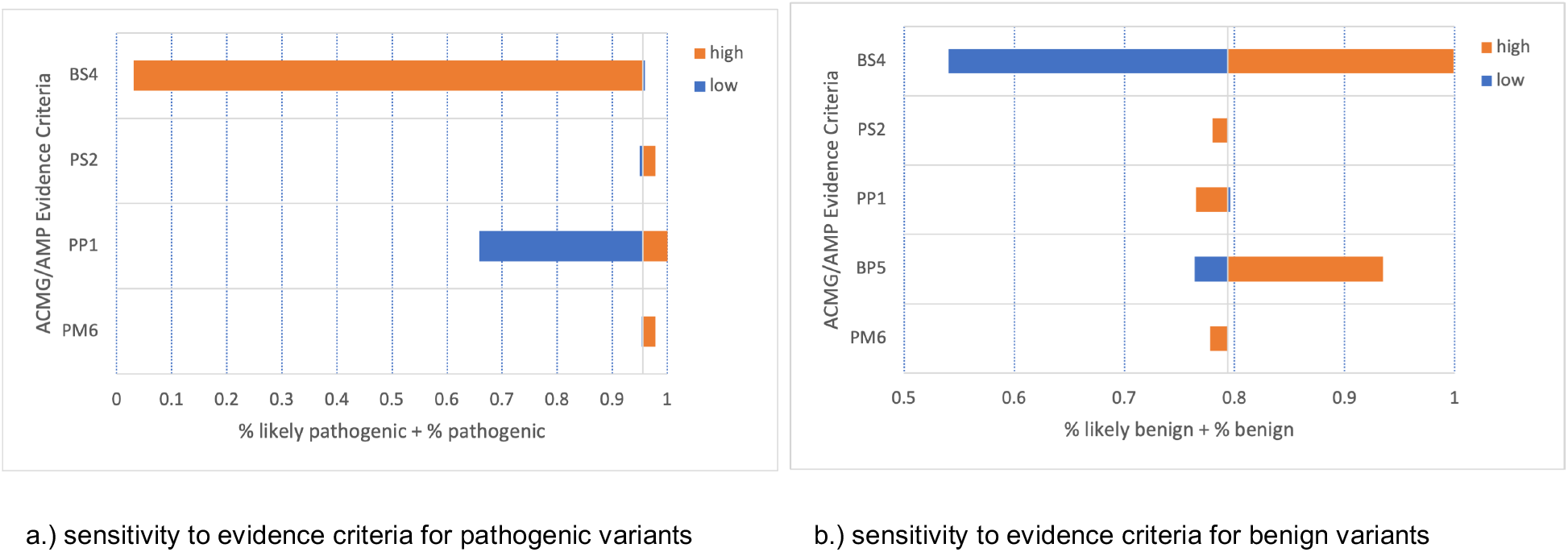
a) Tornado plot for Pathogenic and Likely Pathogenic variants. b) Tornado plot for Benign and Likely Benign variants.

### Probabilities of classifying variants at 1e-06 frequency

For comparison, we evaluated the probability of gathering data for a one-in-a-million variant through data sharing. Figure 5 shows the probability of classifying a 1e-06 variant over the course of 5 years.

**Figure 5:**
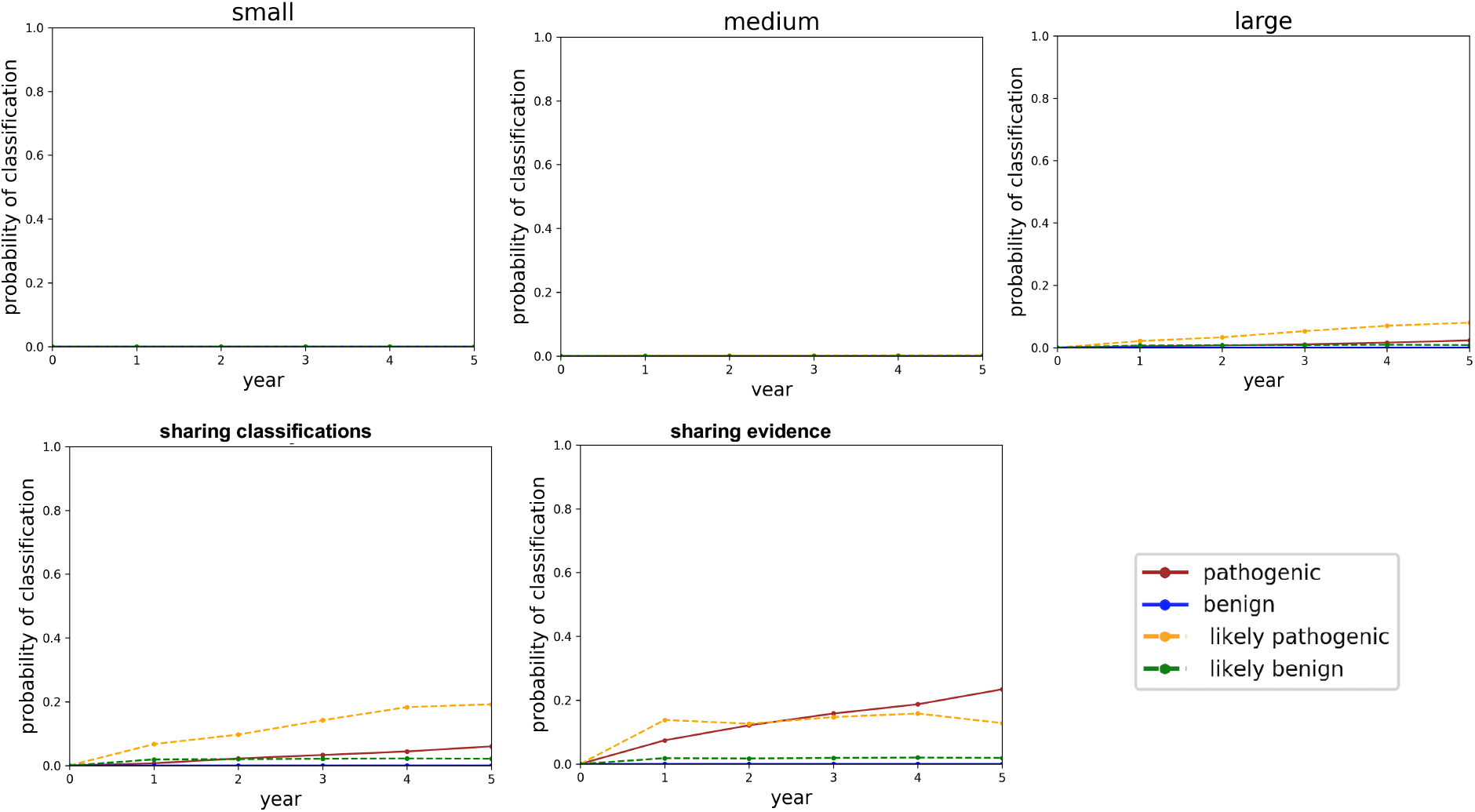
Probabilities of classifying variants at 1e-06 frequency plotted over the course of 5 years. The y-axis of these plots is the probability of classifying the variant, converted from the aggregated likelihoods of pathogenicity generated in the simulations.

In addition to these probability plots, we also performed analysis of 1e-06 variants to generate cumulative odds histograms (supplemental Figure S2) and classification trajectories (supplemental Figure S3). These illustrate similar results. To further explore variant classification timelines for 1e-06 variants, we evaluated classification over 20 years of data sharing (supplemental Figures S4-S6). Sharing evidence is predicted to help classify a minority of 1e-06 variants even after 20 years of data sharing.

## DISCUSSION

These simulations illustrate that clinical data sharing reduces the time and increases the certainty in classifying VUS. Sharing only variant interpretations rather than clinical data, however, results in longer timelines and lower certainty. For example, the same variant could be interpreted as Likely Pathogenic at one laboratory and as a VUS at a different laboratory based on evidence seen at the two respective laboratories. Similarly, the simulations show that evidence for a given variant can, at times, be contradictory. As defined in the ACMG/AMP classification standards, evidence of pathogenicity may be presented for benign variants (and vice versa), though less frequently than for pathogenic variants. Importantly, our simulations demonstrate that discordant evidence resolves more quickly and with higher certainty when centers share their clinical data rather than only sharing their variant interpretations. These are critical results: mis-classified variants mis-inform healthcare providers and may lead to disastrous patient outcomes. [19] Variants originally classified as Likely Pathogenic or Likely Benign more readily become classified as Pathogenic and Benign, respectively, when data is shared.

The simulations also show that classifying pathogenic variants has a higher probability and quicker timeline than for classifying benign variants. The ACMG/AMP evidence criteria and classification guidelines require more evidence for benign classification [7] which results in longer timelines. Models indicate that improved guidelines could balance pathogenic or benign evidence categories, or alternatively create a new “lack of pathogenic evidence despite sufficient observations” category of benign evidence.

We see that highly rare variants (one-in-a-million or less) may be unlikely to be classified by aggregating clinical information alone. Because most variants are highly rare, [20] it’s essential that we invest in strategies for the interpretation of highly rare VUS. One strategy is investment in cascade testing for highly-rare variants in high-penetrance genes. This is an effective strategy because the variant may be rare in the general population but can still be enriched in the family. Another effective strategy is investment in large-scale functional assays, such as MAVEs (Multiplexed Assays of Variant Effect), which can assay thousands of variants at once. [21]

With sufficient clinical data from cooperating sequencing laboratories, these estimates enumerate tangible outcomes that may result from data sharing. There are several mature privacy mechanisms that may be leveraged to share data responsibly; differential privacy, secure multi-party computation, homomorphic encryption, blockchain, and federated computing are approaches that have matured and are available today to protect the privacy of those individuals who have shared their data as well as protect the business interests of the institutions which own the data. [22-25] Additionally, our model can guide functional assay developers as to which variants they should include in their panels. Functional assays are expensive and require expert interpretation, and this information can maximize the impact of those efforts by identifying variant frequencies and sharing scenarios in which data sharing by itself is insufficient for classification.

Most importantly, variant classification timelines will guide prevention, diagnosis, and treatment decisions for patients and their healthcare teams. For example, a patient with a known pathogenic variant in *BRCA1* or *BRCA2* may elect to have a prophylactic mastectomy which, according to the National Cancer Institute, reduces the risk of breast cancer in women who carry a pathogenic *BRCA1* variant by 95%. [26] A patient with a *BRCA1* VUS, on the other hand, may choose to wait if their variant is likely to be classified in the near-term (e.g. within 2 years) but seek alternative options, such as family co-segregation analysis, if that variant will not likely get classified for another 10 years or more. More than half of the variants in the *BRCA1* and *BRCA2* genes are VUS, even though these are two of the most widely studied genes in the human genome.

Other Mendelian diseases with highly penetrant alleles have a significantly larger proportion of VUS, so understanding timelines and probabilities of variant classification will have an even higher impact for those genes.

## CONCLUSION

It is assumed that sharing data should improve variant interpretation. Our research provides a framework to explicitly quantify how much and under what circumstances it improves. We have built and made available a model that simulates the generation and sharing of clinical evidence over time. The software provides graphical results to compare sharing clinical data with sharing only interpretations and sharing nothing. Our experiments were based on data estimates from the literature and from our own experience, but readers can define their own values for the frequencies of observations of various ACMG/AMP evidence criteria as well as experiment with different combinations of centers, different sizes and testing rates, and with different allele frequencies.

## ACKNOWLEDGEMENTS

The authors are very grateful to David Goldgar and Sean Tavtigian for providing their expert insight in the approximation of ACMG/AMP evidence criteria observation frequencies.

