## supplemental text for "Modeling the impact of data sharing on variant classification"

### SUPPLEMENT

In this section, we provide more detail for how we combine evidence in the Bayesian framework. We also provide the graphs for additional experiments run with different configurations, as described in the main text.

### Combining evidence

A single piece of evidence is represented as an odds of pathogenicity. Clinical evidence observed for the same variant from unrelated patients is modeled as independent, so the odds from multiple observations may be combined multiplicatively. The odds of a variant $V_i$ being pathogenic (belonging to the class P) given all the evidence $X_j$ is the product of all the evidence, expressed as odds, as shown in Equation S1.

$$\text{odds}(Vi \in P \mid Xj) = \prod_{j=1}^{n} X_j \qquad \textbf{Equation S1}$$

We convert the odds of pathogenicity to a log scale as shown in Equation S2.

$$\log(\text{odds}(Vi \in P \mid Xj) = \sum_{j=1}^{n} \log(X_j) \qquad \textbf{Equation S2}$$

For a single variant, we compare this sum to the thresholds for Benign, Likely Benign, Likely Pathogenic, and Pathogenic in log scale. The same logic is applied to calculate the odds that the variant is Benign (belonging to the class B), as shown in Equation S3.

$$\log(\text{odds}(Vi \in B \mid Xj) = \sum_{j=1}^{n} \log(X_j) \qquad \textbf{Equation S3}$$

### 10 small, 7 medium, and 3 large centers for 1e-05 over 20 years

The following plots show the same collection of sequencing centers we used in the main text for an allele frequency of 1e-05 but for a timespan of 20 years instead of 5 years.  These plots show that after 20 years of sharing either data or classifications, all the pathogenic variants and almost all benign variants get classified. The difference between the sharing models is how quickly variants are classified and “promoted” from “Likely Benign” and “Likely Pathogenic” to “Benign” and “Pathogenic”, respectively.

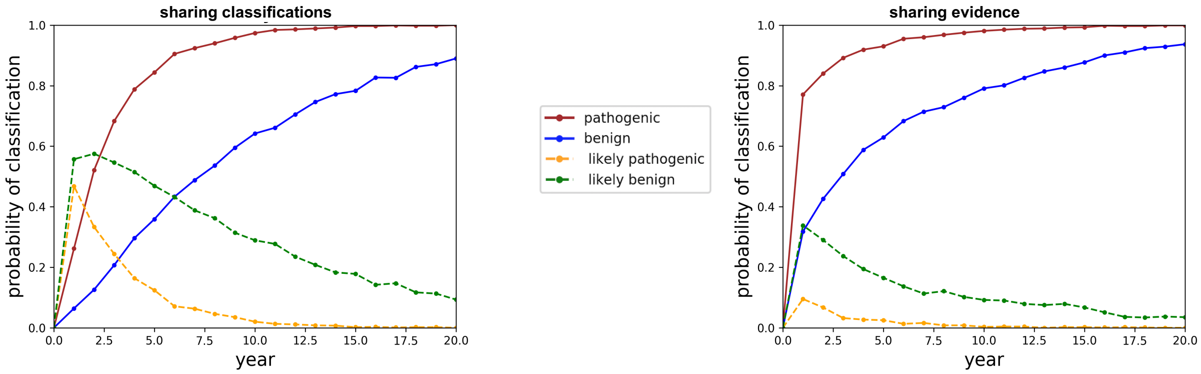

**Figure S1.** Probabilities of classifying variants at 1e-05 frequency plotted over the course of 20 years at 10 small, 5 medium, and 3 large sequencing centers. On the left, all variant classifications and none of the clinical data are shared. On the right, all the clinical data are shared.

<u>10 small, 7 medium, and 3 large centers for 1e-06 over 5 years</u>

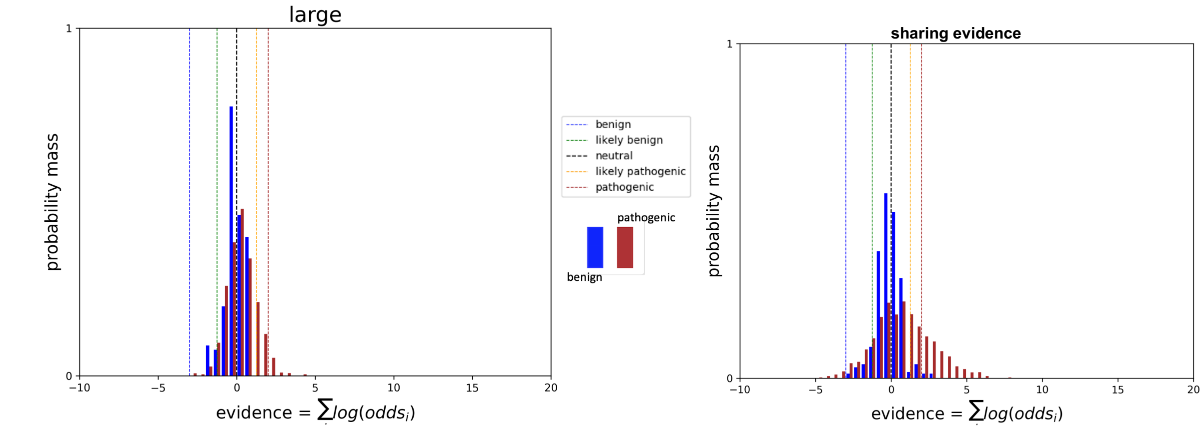


**Figure S2.** Histograms of cumulative log odds for classifying each of 1000 simulated variants present at a 1e-06 frequency in the population for one large sequencing center on the left and for all participating sequencing centers on the right over the course of 5 years. Classification thresholds are demarcated as vertical hash lines. Benign variants are in blue and pathogenic variants in red.

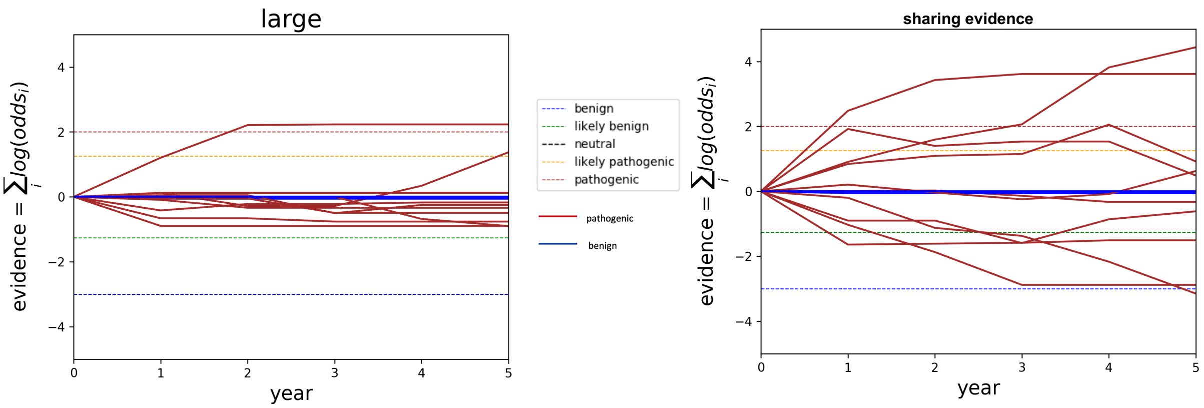


**Figure S3**. Classification trajectories over the course of 5 years for 10 randomly selected variants at 1e-06 frequency in the population for one large sequencing center on the left and for all participating sequencing centers on the right. Classification thresholds are demarcated as horizontal hash lines in the timeline plots. Benign variants are in blue and pathogenic variants in red.

<u>**10 small, 7 medium, and 3 large centers for 1e-06 over 20 years**</u>

The following plots show the same collection of sequencing centers we used in the main text for an allele frequency of 1e-06 but for a timespan of 20 years instead of 5 years.  These plots show that even after 20 years of sharing data, there remains insufficient clinical data to classify variants which occur at 1e-06 frequency in the population.

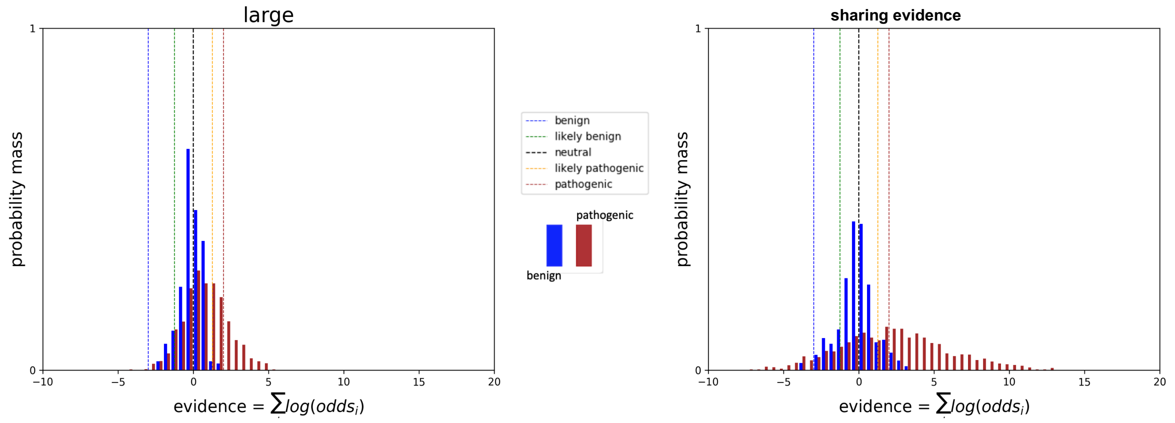


**Figure S4.** Histograms of cumulative log odds for classifying each of 1000 simulated variants present at a 1e-06 frequency in the population for one large sequencing center on the left and for all participating sequencing centers on the right. Classification thresholds are demarcated as vertical hash lines.  Benign variants are in blue and pathogenic variants in red.

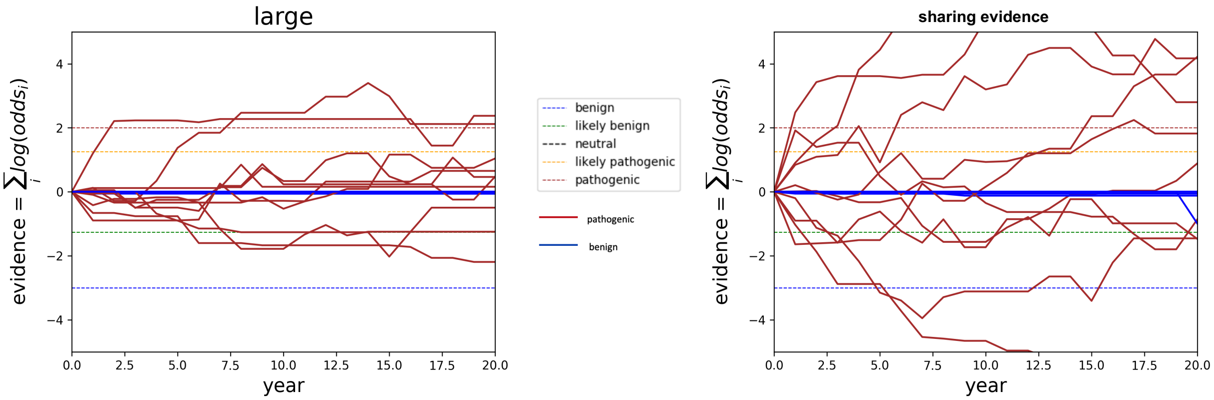


**Figure S5:** Classification trajectories over the course of 20 years for 10 randomly selected variants at 1e-06 frequency in the population for one large sequencing center on the left and for all participating sequencing centers on the right. Classification thresholds are demarcated as horizontal hash lines in the timeline plots. Benign variants are in blue and pathogenic variants in red.

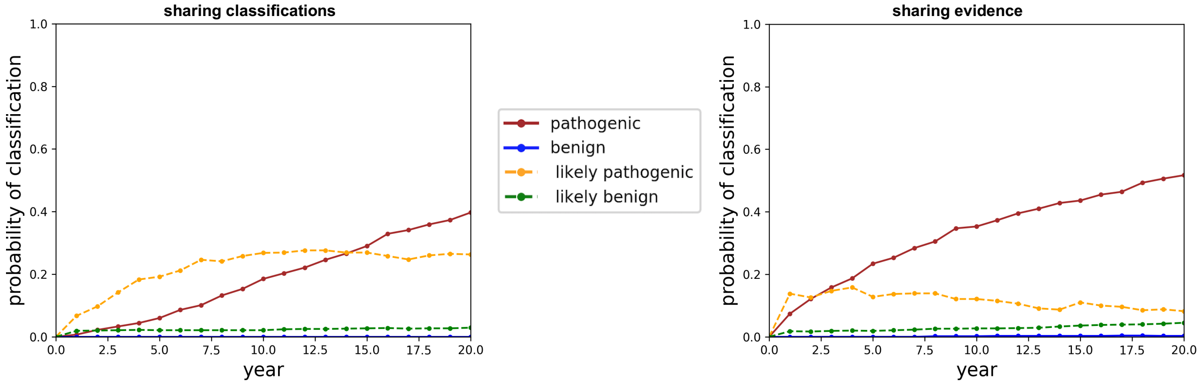


**Figure S6.** Probabilities of classifying variants at 1e-06 frequency plotted over the course of 20 years at 10 small, 7 medium, and 3 large sequencing centers. On the left, all variant classifications and none of the clinical data are shared.  On the right, all the clinical data are shared.

**5 small, 3 medium, and 1 large centers for 1e-5 over 5 years**

The following plots show the same allele frequency and timespan we used in the main text but with a smaller collection of participating centers.  These plots show that sharing clinical data with few centers significantly reduces the probability of classifying variants at the same frequency. The probability of classifying variants as pathogenic reduces from about 95% to about 60%, and the probability of classifying variants as benign reduces from about 60% to about 20%.

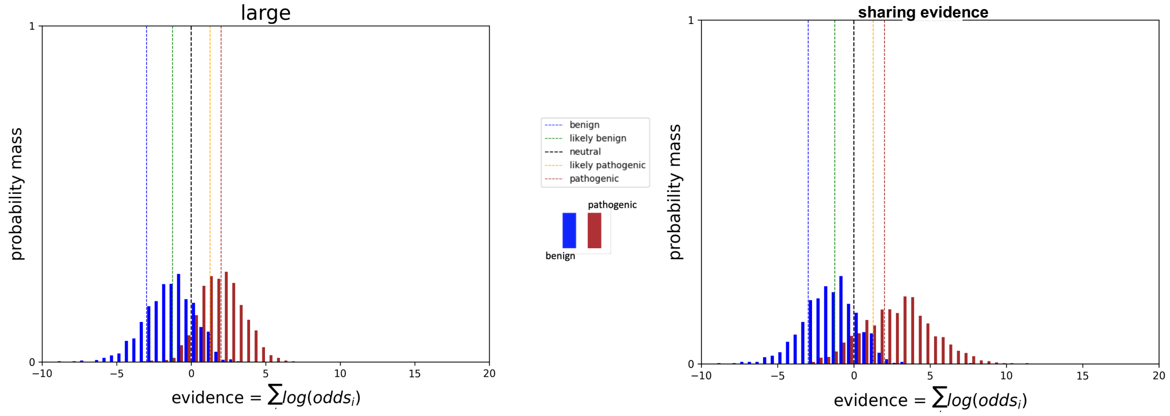


**Figure S7.** Histograms of cumulative log odds for classifying each of 1000 simulated variants present at a 1e-05 frequency in the population for one large sequencing center on the left and for all participating sequencing centers on the right. Classification thresholds are demarcated as vertical hash lines.  Benign variants are in blue and pathogenic variants in red.

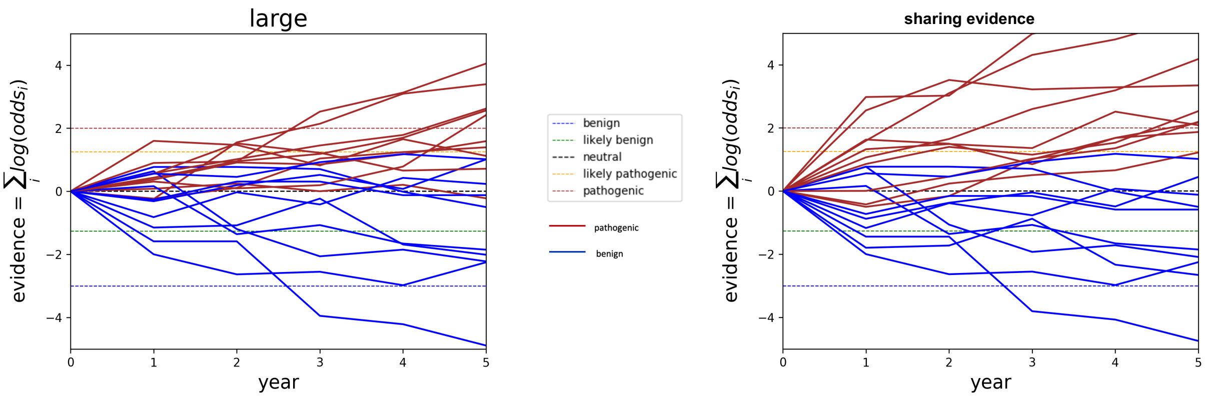


**Figure S8:** Classification trajectories over the course of 5 years for 10 randomly selected variants at 1e-05 frequency in the population for one large sequencing center on the left and for all participating sequencing centers on the right. Classification thresholds are demarcated as horizontal hash lines in the timeline plots. Benign variants are in blue and pathogenic variants in red.

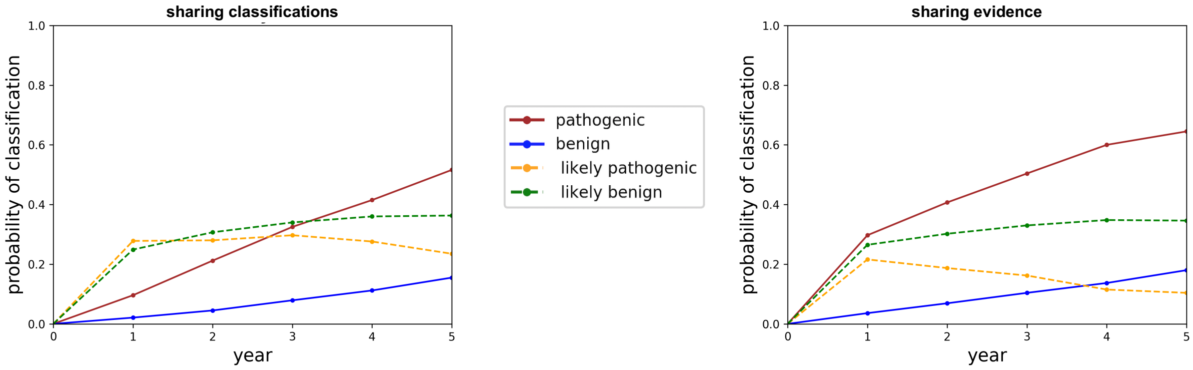


**Figure S9.** Probabilities of classifying variants at 1e-05 frequency plotted over the course of 5 years at 5 small, 3 medium, and 1 large sequencing center. On the left, all variant interpretations and none of the clinical data are shared. On the right, all the clinical data are shared.
